# pydreg: a fast Python package for identifying active *cis*-regulatory elements from nascent transcription

**DOI:** 10.64898/2026.09.06.745329

**Authors:** Adam Y. He, Charles G. Danko

## Abstract

Active promoters and enhancers generate characteristic patterns of RNA transcription that can be measured through nascent RNA sequencing. dREG is a leading method that uses these patterns to identify active *cis*-regulatory elements across the genome, allowing regulatory activity and gene transcription to be profiled in the same experiment. However, its reference implementation was developed around an R-based workflow and a legacy GPU-accelerated support vector machine library that have become increasingly difficult to maintain and deploy. To improve future usability of dREG, we developed pydreg, a Python port of dREG. pydreg preserves the original pretrained models and peak calling procedure from dREG while using contemporary numerical libraries for CPU and GPU computation, resulting in nearly identical peak calls and 5.17 and 5.67-fold reductions in runtime and peak host memory, respectively, compared to dREG. pydreg reduces practical barriers to running dREG locally, improves runtime and memory usage, integrates readily with Python-based genomics workflows, and provides a maintainable foundation on modern computing infrastructure.

## 1 Introduction

*Cis*-regulatory elements (CREs), including enhancers and promoters, coordinate cell-type-specific gene regulation by integrating transcription factor (TF) binding, chromatin state, and 3D genome architecture. A defining feature of many active CREs in mammalian genomes is bidirectional transcription [1, 2], which can be captured by nascent RNA sequencing assays such as GRO-seq [3], PRO-seq [4], and ChRO-seq [5]).

dREG (discriminative regulatory-element detection from GRO-seq) was developed to recognize the characteristic patterns of nascent transcription surrounding active CREs [6]. Rather than relying on a single signal threshold, it summarizes read density at several spatial scales and uses support vector regression (SVR) to identify transcription initiation regions. A subsequent GPU-accelerated version improved sensitivity, resolution, and statistical peak calling [7].

dREG peak calls overlap extensively with orthogonal evidence such as accessible chromatin, H3K27ac, and TF binding, while also recovering weakly active elements that cannot be reliably identified using ChIP-based assays with higher background noise [6, 7]. Even over a decade after its original publication and over 7 years since the publication of its GPU-accelerated successor, dREG remains one of the most accurate methods for identifying active CREs from nascent transcription data [8].

Unfortunately, the reference implementation of dREG is quite complex, combining R, compiled C code, shell scripts, and several external genomics utilities. Its principal GPU accelerator, Rgtsvm, requires a source build against CUDA and Boost [9]. These complex requirements and the legacy nature of dREG and several of its critical dependencies (> 5 years since last update) make deployment and continued maintenance of dREG challenging. Moreover, Rgtsvm uses a custom CUDA implementation designed around early-generation NVIDIA GPUs and sparse workload assumptions that limit its ability to benefit from advances in GPU hardware and software.

To address these usability and performance issues, we developed pydreg, a Python port of dREG. pydreg implements highly optimized I/O, statistical, and machine learning modules; exposes simple command line and programmatic interfaces in a pip-installable package; and substantially improves the computational efficiency, usability, and maintainability of dREG.

## 2 Results

### 2.1 pydreg pipeline overview

pydreg implements the most recent dREG pipeline from [7] to process strand-specific nascent transcription coverage bigWig files into final regulatory element calls (Fig. 1a). It first identifies genomic positions with sufficient read coverage, then summarizes the local read coverage profiles around candidate positions into a single feature vector representing multiple spatial scales. The original pretrained SVR model then scores all feature vectors. Nearby high-scoring positions are assembled into candidate peaks, refined with the original pretrained random forest peak splitting model, assigned statistical significance, and filtered for multiple testing correction.

**Figure 1:**
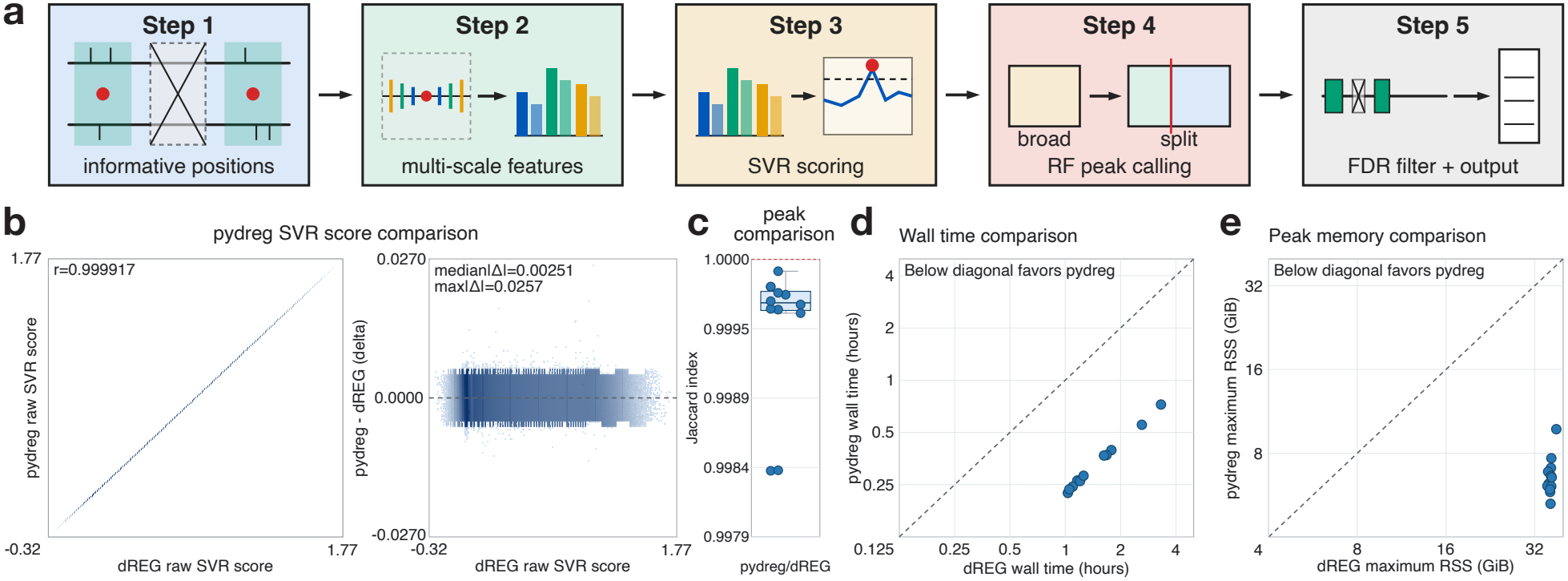
pydreg pipeline, performance, and validation against dREG. (a) Schematic overview of the pydreg/dREG pipeline. (b) Comparison of pydreg and dREG SVR scores (left) and residuals (right) at *n*=126,456,948 candidate positions across 12 libraries. (c) Jaccard index between pydreg and dREG peak calls. (d) Wall time of pydreg and dREG (log scale). (e) Peak host memory of pydreg and dREG (log scale).

The pretrained model parameters distributed with dREG were converted directly to portable safetensors files that are automatically downloaded and cached upon first invocation. pydreg implements a command-line interface to support routine and high throughput analyses, while a Python interface allows pydreg peak calling to be incorporated directly into larger Python-based workflows. Outputs include indexed BED files and bigWig tracks for raw model scores and significant peaks.

### 2.2 Validation and performance

We evaluated pydreg’s outputs and compute efficiency against dREG on an NVIDIA Titan Xp GPU with 16 CPU threads (2*×* Intel Xeon E5 2620 v4) and using 12 previously published GRO/PRO/ChRO-seq datasets (Supplementary Table S1), [1, 5, 7, 10, 11]. pydreg’s SVR implementation produced nearly identical scores compared to dREG (Pearson’s *r* > 0.999, median absolute difference = 0.00251, Fig. 1b). Final peak calls agreed with Jaccard index > 0.998 (median 0.99967, Fig. 1c). pydreg completed runs 4.88 to 5.38 times faster than dREG (median 5.17-fold, Fig. 1d) while reducing maximum resident set size (RSS, host memory) by 3.89 to 6.89-fold (median 5.67-fold, Fig. 1e).

pydreg also runs efficiently on newer NVIDIA and Apple hardware (Supplementary Table S2). On one library (Jurkat_ChROseq_1), four NVIDIA GPUs spanning three architecture generations completed runs in 5.3 to 13.7 minutes, compared to 72 minutes for dREG (benchmarked only on the slowest Titan Xp system). Peak calls remained nearly identical to the official release (Jaccard index 0.99979). Using the MLX backend, an Apple M5 laptop completed the same library in 27.2 minutes with nearly identical peak calls (Jaccard index 0.99951) and the lowest peak host memory of any system tested (4.9 GiB). Optional TensorFloat-32 (TF32) matrix multiplication, available on recent NVIDIA and Apple GPUs, further reduced runtime by 1.21 to 1.51-fold at a small cost in agreement with dREG (Jaccard index 0.996 on NVIDIA and 0.995 on M5).

### 2.3 SVR inference optimization

The SVR at the heart of dREG is a standard radial basis function (RBF, Gaussian)-kernel SVR that operates on a length 360 feature vector with 605,187 support vectors. As RBF SVR inference times scale linearly with respect to the product of the number of support vectors and the feature dimension, prediction with the dREG SVR is quite computationally intensive.

The original authors of dREG eventually implemented GPU acceleration using their own custom library Rgtsvm [9]. Rgtsvm modifies the earlier GTSVM library [12], which was developed for early CUDA hardware and sparse datasets. During prediction, pairwise kernel values are evaluated using custom sparse-oriented CUDA kernels rather than by expressing dense query–support vector dot products as chunked matrix multiplications. For dREG’s dense 360-dimensional features, this organization provides less data reuse than general matrix multiplication (GEMM) and can increase device memory traffic.

To address these performance limitations, we reimplemented dREG in Python, whose GPGPU ecosystem is broader and more mature than R’s [13]. pydreg uses CuPy [14] and MLX [15] for GPU-accelerated SVR scoring on NVIDIA and Apple silicon GPUs, respectively, with an efficient CPU fallback implemented using NumPy and Numba [16, 17].

During SVR scoring, query batches *X* ∈ ℝ^*Q×d*^ are evaluated against support vector chunks 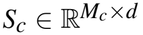. We use the identity

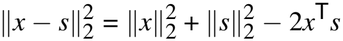

to calculate pairwise squared distances from row norms and the matrix product 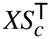. A fused elementwise kernel combines the matrix product with the row norms and applies the RBF transformation, after which a matrix vector product weights the SVR kernel values by the dual coefficients. This formulation enables the dominant computations in SVR prediction to be calculated using vendor-optimized GEMM kernels. Precomputed row norms and fused kernels further reduce redundant arithmetic and device memory traffic.

To test the efficiency of our new GPU-accelerated scoring implementation, we used Nsight Systems (version 2025.3.1) to profile GPU utilization, kernel launches, and host-to-device (H2D) memory copies during the primary GPU-scoring step of dREG and pydreg. We found that, using default settings, pydreg performed 12*×* and 942*×* as much work per kernel launch and H2D memory copy, respectively, as dREG (median, Supplementary Fig. S1). Total scoring was 6.75*×* faster using pydreg (Supplementary Fig. S2). We note that dREG has substantially lower volatile GPU utilization (67.4% vs 95.4%, Supplementary Fig. S3). This is due to inefficient I/O and feature extraction functions that are run serially in between batched GPU calculations; pydreg significantly optimizes these steps and performs prefetching, which substantially reduce CPU bottlenecks during GPU scoring (Supplementary Fig. S2). Nonetheless, GPU-active compute time is still 5.47*×* shorter in pydreg due to the aforementioned GPU optimizations (median, Supplementary Fig. S2).

### 2.4 Statistical peak calling optimization

Apart from SVR scoring, the most computationally demanding portion of dREG’s algorithm is false discovery rate (FDR)-controlled peak calling. For each candidate summit, dREG selects five smoothed local scores **x** ∈ ℝ^5^ and computes

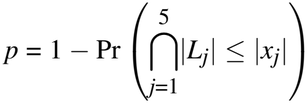

under a correlated five-dimensional Laplace null

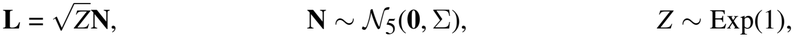

where Σ ∈ ℝ^5*×*5^ is a covariance matrix calculated once per dataset that represents expected background correlation across nearby scored loci. The integral over *Z* is evaluated on a fixed grid, with randomized quasi-Monte Carlo (QMC) used at each retained grid point to estimate the corresponding five-dimensional Gaussian box probability. dREG then reports peak calls with Benjamini-Hochberg-adjusted *p* ≤ 0.05.

pydreg skips zero-weight grid points and terminates integration once the remaining tail’s maximum possible contribution falls below 10^−6^, eliminating unnecessary QMC evaluations with either zero or provably bounded effect. The former optimization is mathematically exact, whereas the latter was found empirically to lead to no practical degradation in statistical calculations. Collectively, these optimizations reduce QMC evaluations by about half, substantially accelerating *p*-value calculation.

## 3 Conclusion

By preserving the published dREG algorithm while adopting modern numerical libraries and packaging, pydreg extends the useful life of dREG and improves its reproducibility and interoperability with current workflows. We also dramatically reduce the computational resources needed to run dREG.

## 4 Conflicts of interest

The authors declare that they have no competing interests.

## 5 Funding

This work is supported in part by funds from the Astera Institute (https://ror.org/00ydx1s47) through the 2026 Hidden Science Competition. To comply with the Astera Open Science Policy [18], we decline formal journal publication for this manuscript. The bioRxiv version is the version of record.

## 6 Code and data availability

pydreg is implemented in Python 3.11+ and is freely available under the GPL-3 license at https://github.com/adamyhe/pydreg and from PyPI via `pip install pydreg[gpu]` (for CUDA acceleration) or `pip install pydreg[mlx]` (for Apple Metal acceleration). The version as of the publication of this manuscript is 1.0.0 (https://doi.org/10.5281/zenodo.24512251). Pretrained model parameters are automatically downloaded and cached from HuggingFace (https://doi.org/10.57967/hf/10017).

No new sequencing data were generated in this study. Previously published data and pydreg/dREG outputs underlying the benchmarks in Fig. 1 are available from the Gene Expression Omnibus (Supplementary Table S1) and HuggingFace (https://doi.org/10.57967/hf/10018).

A pre-built Apptainer image for dREG and rdata-formatted model parameters are preserved on Zenodo (https://doi.org/10.5281/zenodo.10113378), with run and build instructions at (https://github.com/Danko-Lab/dREG-apptainer).

## 7 Author contributions statement

A.Y.H. and C.G.D. conceived of the study. A.Y.H. implemented the code and performed benchmarking experiments with guidance from C.G.D. A.Y.H. wrote the manuscript with revisions from C.G.D.

## 8 Generative AI use disclosure

Coding agents (Claude Code running Opus 4.6, 5, and 5.5 and Sonnet 5 and Codex running GPT-5.5) were used to write the code base underlying this manuscript.

## 9 Acknowledgments

The authors thank the users of dREG, whose many bug reports motivated the development of pydreg. The authors also thank the previous developers of dREG, upon whose work pydreg is built.

Some of the computing for this project was performed on the Sherlock cluster at Stanford University. We would like to thank Stanford University and the Stanford Research Computing Center for providing computational resources and support that contributed to these research results.

---

**Supplementary Table S1:** GRO/PRO/ChRO-seq libraries used in the pydreg-vs-dREG benchmarks.

| Internal library ID | GEO accession | Biosample | Assay | Ref |
|---|---|---|---|---|
| G1 | GSM1480327 | K562 | PRO-seq | [1] |
| G3 | GSM3452725 | K562 | PRO-seq | [7] |
| G5 | GSE89230 | K562 | PRO-seq | [11] |
| G6 | GSM2545324 | K562 | PRO-seq | [10] |
| G7 | GSM2545325 | K562 | PRO-seq | [10] |
| GM12878_groseq | GSM1480326 | GM12878 | GRO-seq | [1] |
| K562_groseq | GSM1480325 | K562 | GRO-seq | [1] |
| Jurkat_PROseq | GSM3309955 | Jurkat | PRO-seq | [5] |
| Jurkat_ChROseq_1 | GSM3309957 | Jurkat | ChRO-seq | [5] |
| Jurkat_ChROseq_2 | GSM3309956 | Jurkat | ChRO-seq | [5] |
| Jurkat_ChROseq | GSM3309956, GSM3309957 | Jurkat | ChRO-seq (pooled replicates) | [5] |
| Jurkat_leChROseq | GSM3309958 | Jurkat | length extension (le)ChRO-seq | [5] |

**Supplementary Table S2:** pydreg performance across hardware on one library (Jurkat_ChROseq_1), with --seed 0. Jaccard index indicates comparison against the dREG results.

|  |  | Wall time (min) |  | Peak RSS | Jaccard index |  |
|---|---|---|---|---|---|---|
| GPU | CPU | float32 | TF32 | (GiB) | float32 | TF32 |
| NVIDIA L40S | Xeon Gold 6426Y | 5.3 | 4.4 (1.21×) | 6.0 | 0.99979 | 0.99618 |
| NVIDIA RTX 3080 | Ryzen 7 9800X3D | 6.3 | 5.0 (1.26×) | 6.2 | 0.99979 | 0.99618 |
| NVIDIA A100 | EPYC 7763 | 7.4 | 4.9 (1.51×) | 6.0 | 0.99979 | 0.99618 |
| NVIDIA Titan Xp | 2× Xeon E5-2620 v4 | 13.7 | n/a | 5.8 | 0.99979 | n/a |
| Apple M5 (MLX) | Apple M5 | 27.2 | 18.1 (1.50×) | 4.9 | 0.99951 | 0.99469 |
| dREG, Titan Xp | 2× Xeon E5-2620 v4 | 72.2 | n/a | 36.1 | n/a | n/a |

**Supplementary Figure S1:**
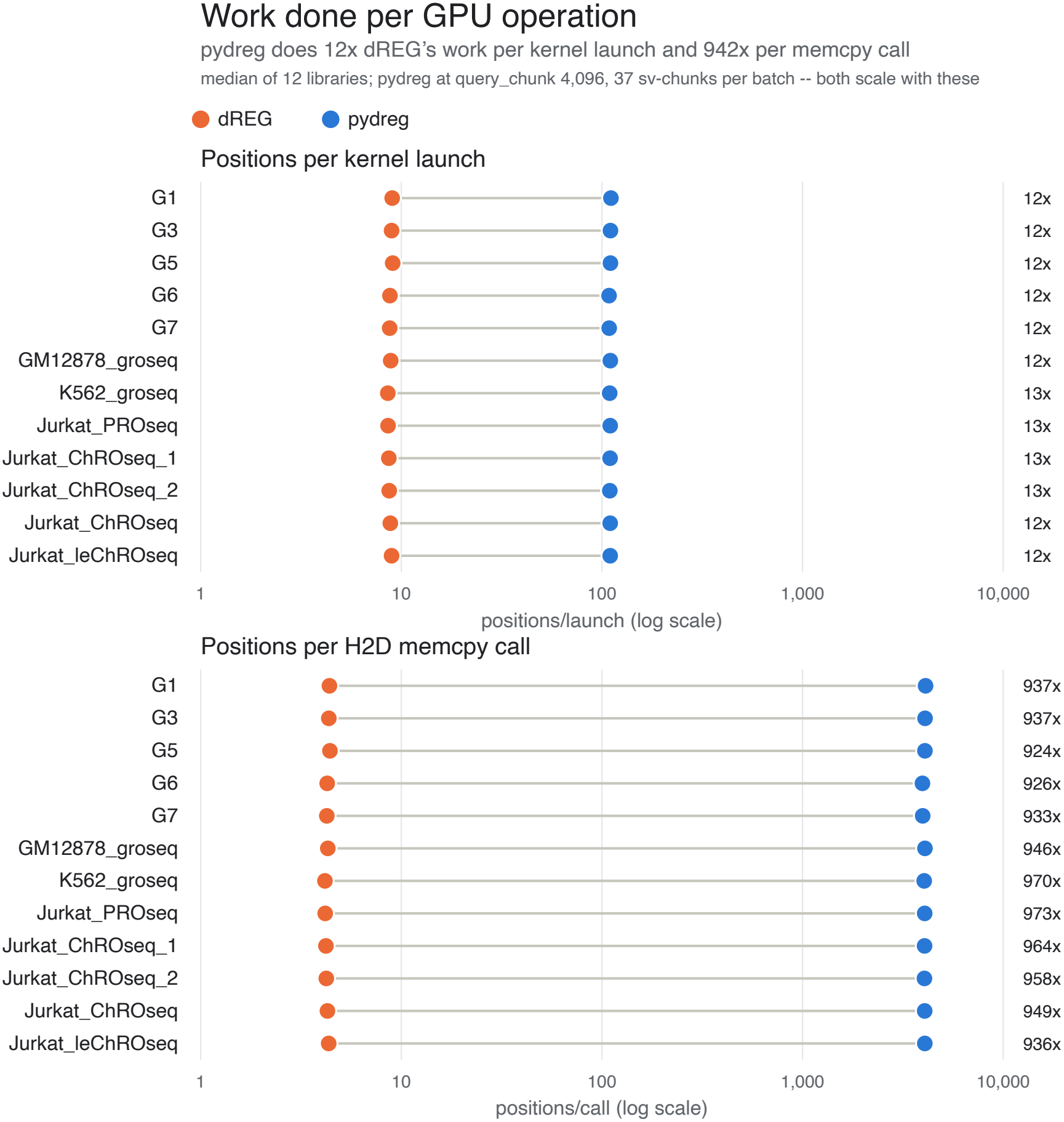
GPU efficiency (amount of work done per kernel launch or H2D memory copy) of dREG and pydreg.

**Supplementary Figure S2:**
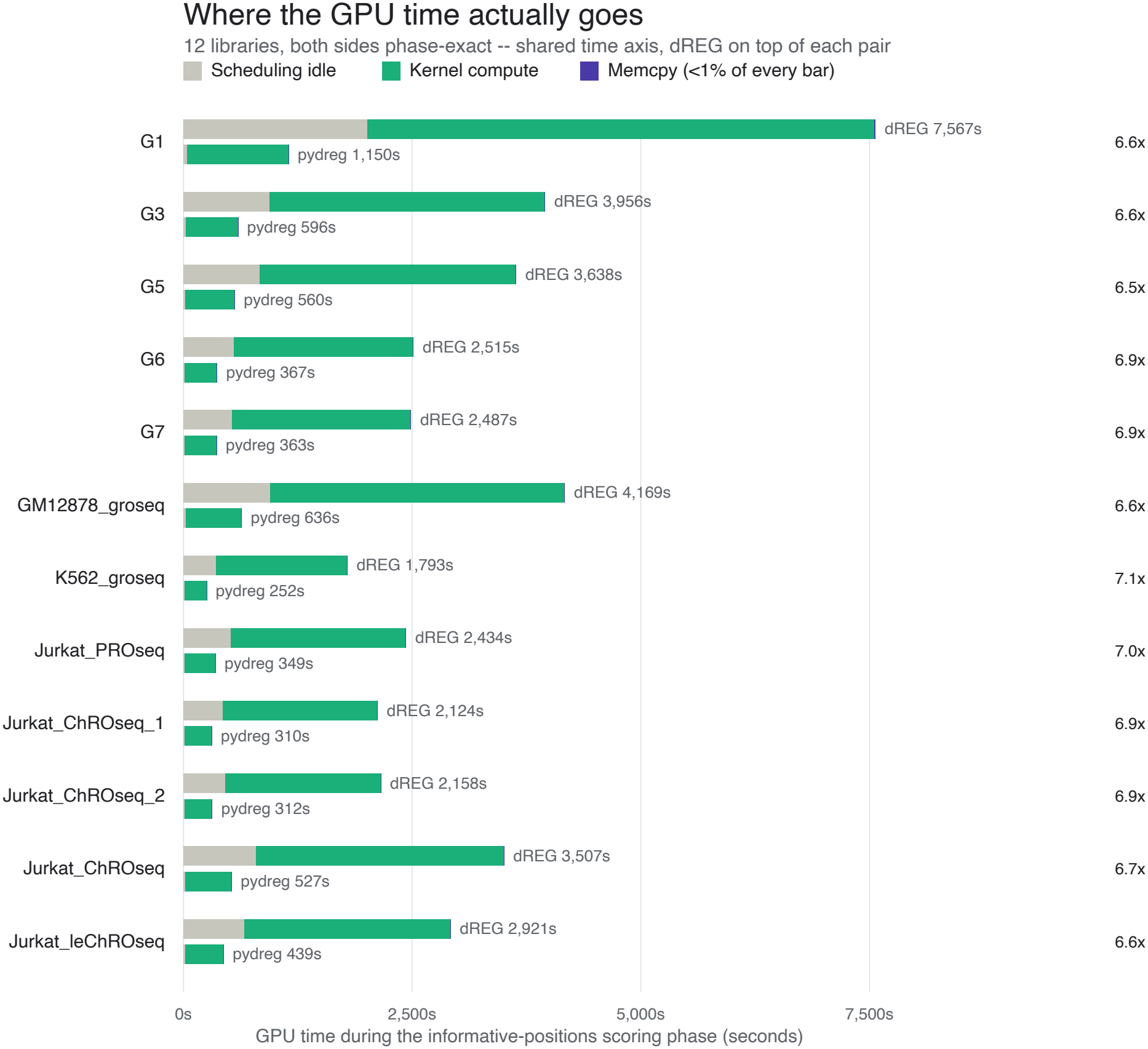
Breakdown of GPU compute stages by wall time for dREG and pydreg.

**Supplementary Figure S3:**
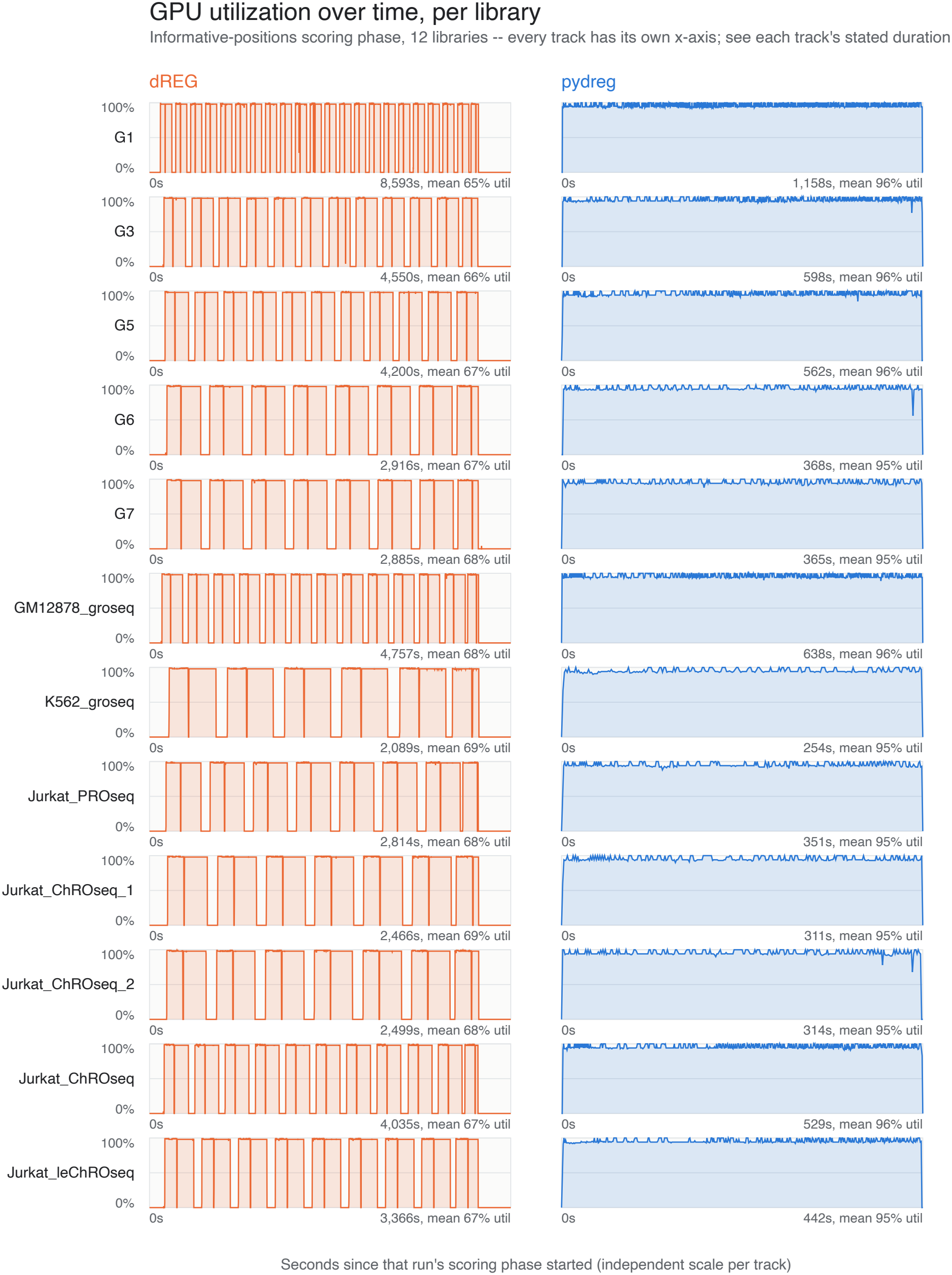
GPU utilization traces for dREG and pydreg.

## Notes

https://github.com/adamyhe/pydreg

https://doi.org/10.57967/hf/10017

https://doi.org/10.57967/hf/10018

### Competing Interest Statement

The authors have declared no competing interest.

### Summary of Updates

We have further optimized the pydreg package and released a stable 1.0.0 version. This revision updates the technical note to describe benchmarking results conducted on v1.0.0.

